# Results of the Protein Engineering Tournament: An Open Science Benchmark for Protein Modeling and Design

**DOI:** 10.1101/2024.08.12.606135

**Authors:** Chase Armer, Hassan Kane, Dana L. Cortade, Henning Redestig, David A. Estell, Adil Yusuf, Nathan Rollins, Hansen Spinner, Debora Marks, TJ Brunette, Peter J. Kelly, Erika DeBenedictis

## Abstract

The grand challenge of protein engineering is the development of computational models to characterize and generate protein sequences for arbitrary functions. Progress is limited by lack of 1) benchmarking opportunities, 2) large protein function datasets, and 3) access to experimental protein characterization. We introduce the Protein Engineering Tournament—a fully-remote competition designed to foster the development and evaluation of computational approaches in protein engineering. The tournament consists of an *in silico* round, predicting biophysical properties from protein sequences, followed by an *in vitro* round where novel protein sequences are designed, expressed and characterized using automated methods. Upon completion, all datasets, experimental protocols, and methods are made publicly available. We detail the structure and outcomes of a pilot Tournament involving seven protein design teams, powered by six multi-objective datasets, with experimental characterization by our partner, International Flavors and Fragrances. Forthcoming Protein Engineering Tournaments aim to mobilize the scientific community towards transparent evaluation of progress in the field.

**Graphical Abstract:** 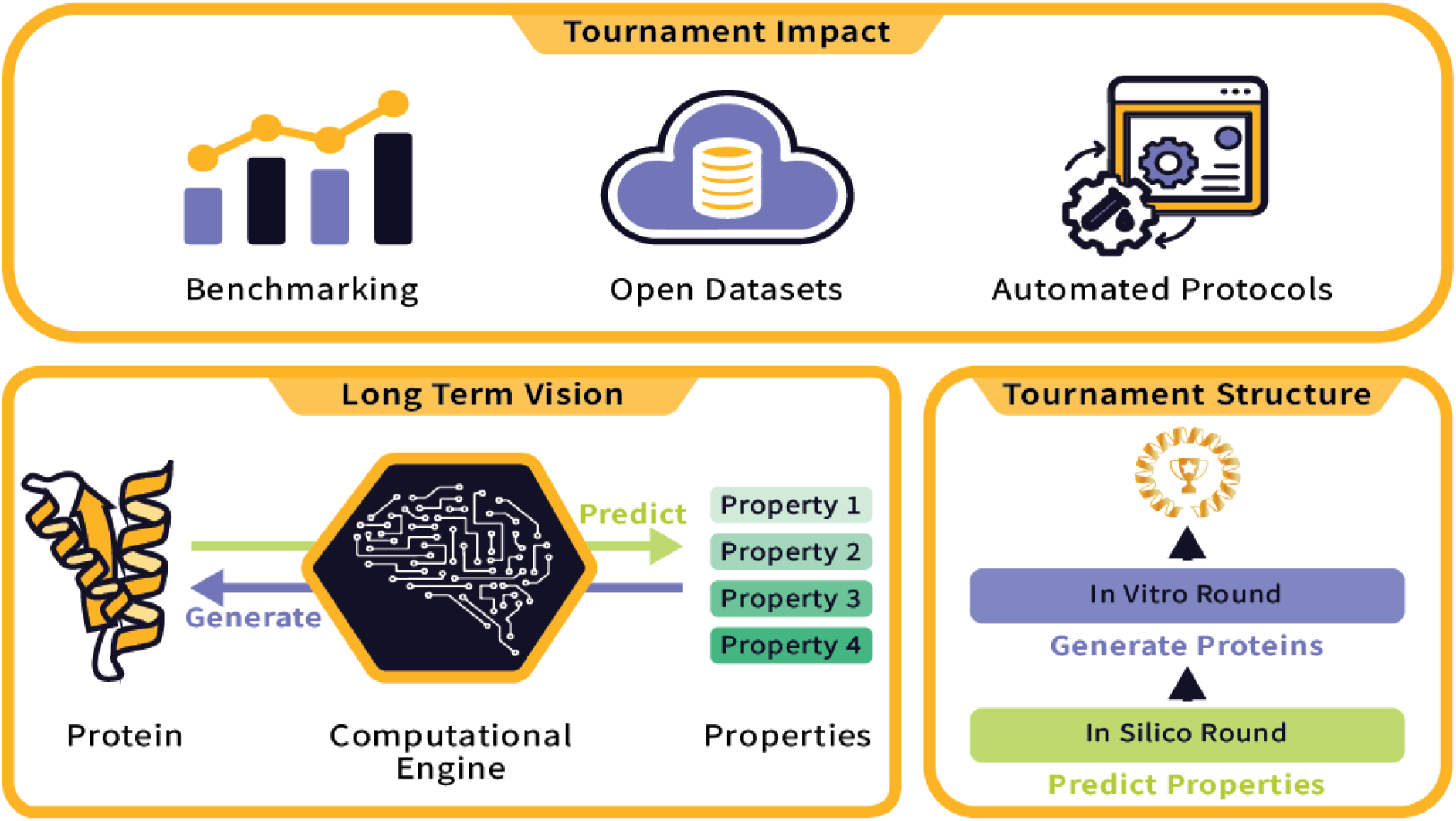

---

The field of computational protein engineering aims to improve our understanding of protein function and design by developing predictive models, which infer biophysical properties from a protein’s sequence^1^, and generative models, which compose protein sequences possessing a desired set of properties^2^. However, several obstacles are currently limiting model development in both paradigms.

Predictive modeling has been notably hampered by the lack of large, complex, and diverse datasets. Public, open datasets have long provided valuable opportunities for developing and benchmarking new methods in machine learning research. Computer vision datasets, such as MNIST^3^ and ImageNet^4^, enabled individual research labs to experiment with new approaches, and also created a benchmark with which to measure collective progress. Researchers in the protein engineering community have also utilized open datasets, such as FLIP^5^ and TAPE^6^, and curated datasets of hundreds of deep mutational scanning experiments like ProteinGym^7^ to encourage similar developments. However, most available datasets, such as those documented in ProtaBank^8^, are limited in scope, predominantly mapping simple sequence-function relationships through single point mutations. This simplicity restricts the ability of predictive models to accurately characterize a wide range of protein functions under varied conditions.

Generative modeling also faces its own set of challenges, primarily due to computational scientists’ limited capacity to experimentally validate and characterize their protein designs. This gap significantly hinders the development and benchmarking of new generative design methods, as there is no standardized evaluation criteria to reliably test and compare the efficacy of these novel protein sequences. Addressing these challenges is crucial for the advancement of computational methods in protein engineering, as it would enable more accurate predictions and innovative designs in protein function.

Science competitions have historically tackled predictive benchmarking efforts by allowing researchers to test their computational methods on never-before-seen datasets. Perhaps the most notable example is the Critical Assessment of Structure Prediction (CASP)^9^ a biennial event for computational protein structure prediction. Since its inception, CASP has become a crucial benchmark for the protein structure prediction community. By creating visibility around a single event, the competition has inspired an ambitious spirit among researchers to develop the best performing method, thereby encouraging rapid development. CASP has inspired the creation of similar competitions, like the Critical Assessment of Computational Hit-finding Experiments (CACHE)^10^, which was created in the computational chemistry field to benchmark novel approaches for finding new small-molecule binders.

We believe there is a burgeoning opportunity to create a new scientific competition that addresses the unique challenges of both predicting and engineering protein function. We previously outlined these challenges and proposed that a Protein Engineering Tournament could tackle the aforementioned obstacles by curating a series of tournaments which provide novel datasets for predictive modeling and experimental validation for generative design^11^. By introducing never-before-seen datasets on protein function and offering open-source experimental characterization of novel proteins, the Protein Engineering Tournament hopes to overcome this barrier and enable diverse research groups from all backgrounds to participate in cutting-edge protein engineering research. The Tournament hopes to act as a unifying force for the community by providing a transparent platform for benchmarking protein modeling and design methods, reducing barriers to model development and validation, and accelerating the field of protein engineering.

## Results

### Tournament structure

To enable benchmarking of both predictive and generative models, the Tournament was structured to have an *in silico* round to benchmark predictive models followed by an *in vitro* round to benchmark generative models (Fig. 1). In the *in silico* round, the teams were tasked with developing models for predicting biophysical properties of given protein sequences (Fig. 1A). In the *in vitro* round (Fig. 1B), the teams were asked to design protein sequences that maximize or satisfy certain biophysical properties. Following the conclusion of the tournament, participating teams were asked to submit abstracts, detailing the methods applied in both the *in silico* and *in vitro* rounds (see Methods).

**Figure 1:**
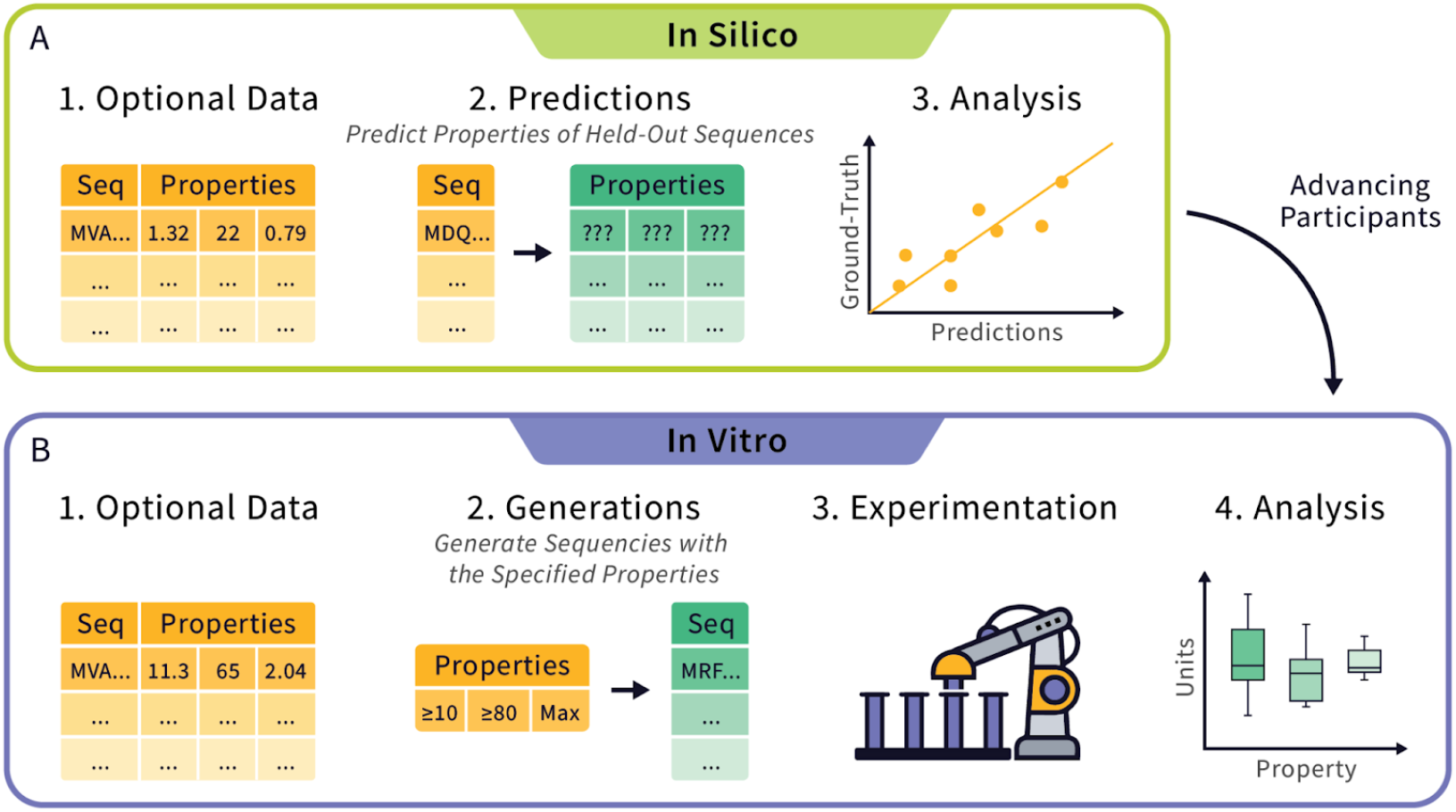
Overview of Tournament Rounds. The tournament consists of two rounds: (A) the *in silico* round, where participants predict properties from protein sequences, with optional training data, and (B) the *in vitro* round, where participants generate protein sequences that possess a desired set of properties which are then to be expressed and characterized. Initial interest in the pilot tournament led to the registration of over 90 individuals assembled into 28 teams, representing a mix of academic (55%), industry (30%), and independent (15%) teams. For the *in silico* round, 7 teams successfully submitted predicted properties (Supp. Table 1). Teams that performed well in at least one *in silico* event were invited to participate in the *in vitro* round (See Methods). The *in vitro* round consisted of 5 teams that advanced from the *in silico* round and 2 generative teams that were invited to participate (Supp. Table 1).

### Datasets and Events

The pilot Protein Engineering Tournament was made possible by the donation of 6 multi-objective datasets from our academic and industry partners (Table 1) structured into two rounds (Fig. 2). Each dataset was used as a unique event in the Tournament, with the exception of the **α**-Amylase dataset which was used in three events. For each **α**-Amylase event, the challenge problem and dataset was modified to reflect the track (zero-shot or supervised) and round *(in silico* and *in vitro*). By using donated datasets, the Tournament provided an opportunity for academic and industry groups to disseminate privately held or previously unpublished datasets to the protein engineering community.

**Table 1:** Datasets used in the pilot Tournament. *The enzyme target and corresponding donor for each dataset are displayed along with the Round (in silico* and/or *in vitro*), Track (zero-shot and/or supervised), and the Challenge Problem posed to the teams. See supplementary Table 2 for suggested citations, links to GitHub, and additional references material for each dataset.

| Enzyme Target | Dataset Details (total data points) | Donor | Round (Track), and “Challenge Problem” |
| --- | --- | --- | --- |
| Aminotransferase | 147 sequences measured against three different substrates (441 data points) | Prof. Dr. U. Dr. U.T. Bornscheuer and M.Sc. M. J. Menke from the University of Greifswald, Dept. of Biotechnology & Enzyme Catalysis | *in silico* (zero-shot):<br>“Score how active you predict each variant is for each of the three substrates (e.g. log probabilities):<br>1. S-Phenylethylamine<br>2. (4-Chlorophenyl)phenylmethanamine<br>3. 1,3-Diphenyl-propane-1-amine.<br>The range of scoring is arbitrary.” |
| α- Amylase | 9422 sequences measured for expression, specific activity, and thermostability (28,266 data points made from 7578 single mutations) | International Flavors and Fragrances (IFF) | *in silico* (zero-shot and supervised):<br>“Score the following three properties for each variant (e.g. log probabilities):<br>1. specific activity<br>2. Expression<br>3. thermostability<br>The range of scoring is arbitrary.” |
|  | and 1843 rationally combined multi-mutants + wildtype) |  | *in vitro*:<br>“Submit a list of up to 200 Amino Acid Sequences that maximize enzyme activity while maintaining 90% stability and expression of the parent sequence. Your list of sequences should be ranked.” |
| Imine Reductase | 4517 sequences measured for activity (4517 data points) | Codexis | *in silico (supervised)*:<br>“Score the fold improvement over positive control (FIOP) of activity. The range of scoring is arbitrary.” |
| (table continued on next page)<br>Alkaline Phosphatase PafA | 1,041 sequences measured against three substrates (3, 123 data points) | The Polly Fordyce Group, Stanford University, with guidance from Dr. Craig Markin and Prof. Polly Fordyce | *in silico (supervised)*:<br>“Score activity for each of the three substrates:<br>1. methyl phosphate (MeP) - chemistry limited substrate<br>2. Carboxy 4-methylumbelliferyl phosphate ester (cMUP) - binding limited substrate<br>3. methyl phosphodiester (MecMUP) - promiscuous substrate<br>The range of scoring is arbitrary.” |
| β-Glucosidase B | 456 sequences measured for activity and melting point (912 data points) | The students and faculty members of the Design to Data (D2D) undergraduate research program, led by Ashley Vater and Professor Justin B. Siegel at University of California, Davis | *in silico (supervised)*:<br>“Score each of the following properties:<br>1. Activity<br>2. Melting point<br>The range of scoring is arbitrary.” |
| Xylanase | 201 sequences measured for expression (201 data points) | The Sarel Fleishman Lab from the Weizmann Institute of Science | *in silico (zero-shot)*:<br>“Given the sequence, please predict how well the enzyme expresses. Your prediction should be a classification (0=No expression, 0.5=Low expression, 1=Good expression). The range of scoring is arbitrary.” |

**Figure 2:**
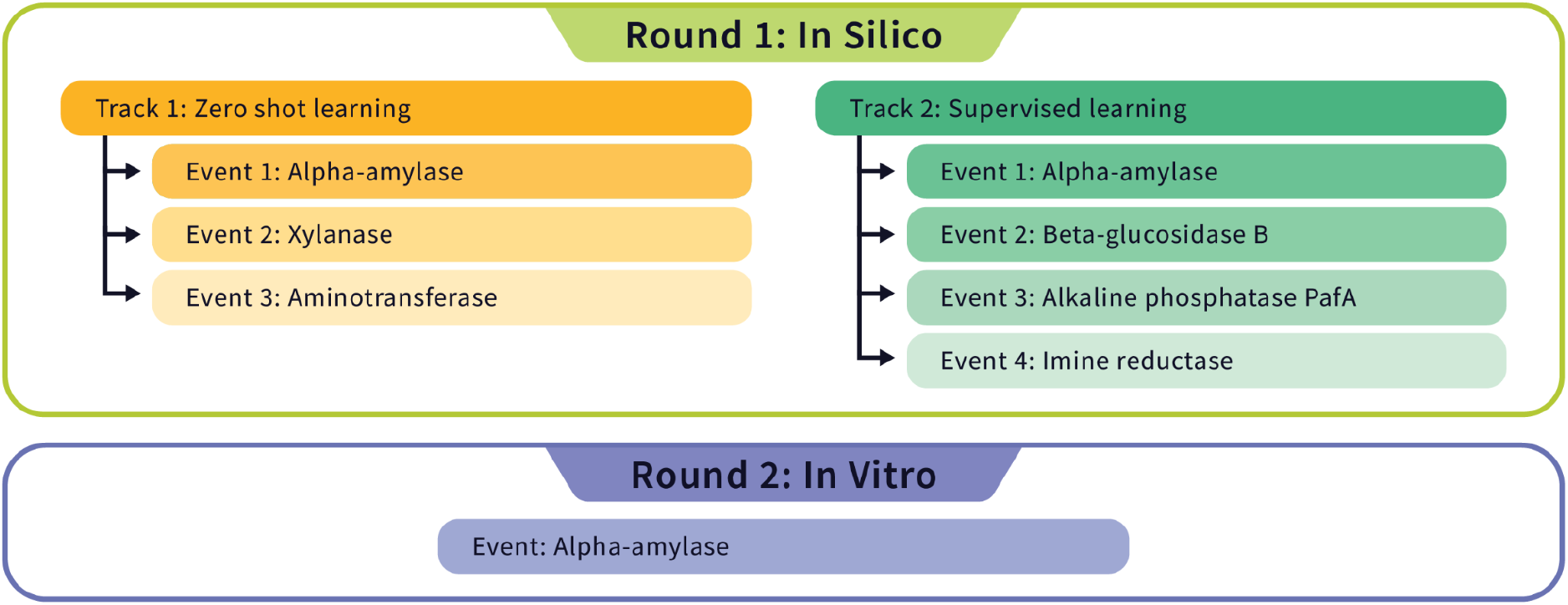
Tournament Structure. Events from the pilot Protein Engineering Tournament organized by Round (*in silico* or *in vitro*) and Track (zero-shot or supervised).

### The *in silico* round

In the *in silico* round participants engaged in seven events distributed across two unique tracks: zero-shot and supervised. Teams had the flexibility to choose their level of participation, opting into any number of these events. The objective for each event was to predict a specified set of properties—such as expression levels, thermostability, and specific activity—from provided protein sequences (Table 1).

The zero-shot track challenged participants to make predictions without any prior training data, relying solely on intrinsic algorithmic robustness and generalizability. This track comprised three events centered around the enzymes α-amylase, aminotransferase, and xylanase. Teams were given three weeks to access the datasets and make submissions.

For the supervised track, participants received a pre-split training and test dataset for each event. The training datasets included sequences and their measured properties, while the test datasets included sequences without measured properties. Participants were directed to train their models using the training set and subsequently use the trained model to predict the withheld properties in the test set. Participant’s test set submissions were assessed against the ground-truth withheld data. This track included four events focusing on different enzymes: alkaline phosphatase PafA, α-amylase, β-glucosidase B, and imine reductase. Teams were given six weeks to access the datasets and make submissions.

For all *in silico* events, teams’ results were used to determine their rank and were subsequently awarded points based on their performance (Fig. 3) (see Methods for details on evaluation metrics and scoring). By summing the points received in each event, leaderboards were generated for both the zero-shot and supervised events (Fig. 3), as well as a combined *in silico* leaderboard (Fig. 3). The Marks Lab won the zero-shot track, while Exazyme and Nimbus shared the first-place spot for the supervised track. Based on combined performance across both tracks, Nimbus was the overall winner for the *in silico* round.

**Figure 3:**
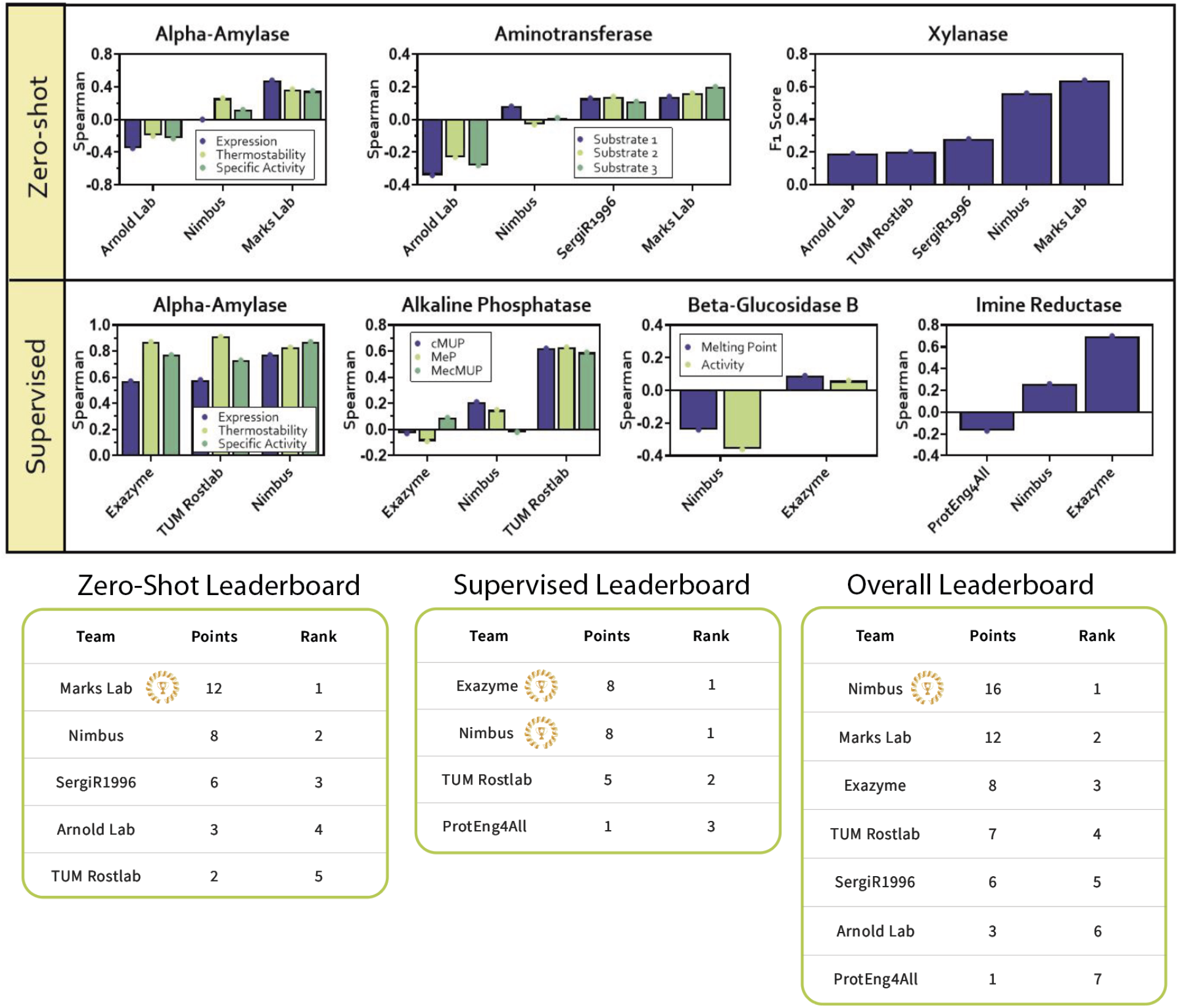
The *in silico* event results. Results for all *in silico* events, including the leaderboards for the Zero-Shot track, Supervised track, and combined overall *in silico* leaderboard. All results are shown as either Spearman (Spearman Correlation), FIOP (Fold-Improvement Over Positive control), or the weighted F1 Score. Definitions for the Alkaline Phosphatase event include: cMUP= Carboxy 4-methylumbelliferyl phosphate ester, MeP= methyl phosphate, and MecMUP= methyl phosphodiester. Important definitions for the Aminotransferase event include: Substrate 1=S-Phenylethylamine, Substrate 2=(4-Chlorophenyl)phenylmethanamine, Substrate 3 =1,3-Diphenyl-propane-1-amine.

### The *in vitro* round

In the *in vitro* round, teams were tasked with using their computational models to propose novel enzyme designs that exhibited enhanced functional properties. Teams were provided with data on single and rationally designed multi-mutations of the **α**-amylase enzyme, up to 11 mutations from wildtype. Teams were instructed to generate a ranked list of up to 200 protein sequences. The criteria for these sequences were multifold: they should maximize enzyme activity and retain at least 90% of the stability and expression levels of the parent enzyme. Not all teams submitted the full 200 sequences, and not all submitted sequences were able to be synthesized, cloned and/or expressed (Supp. Table 3). Teams used a variety of approaches for their generation: MediumBio only used assay-labeled data; TUM used LLMs to further engineer the top performing IFF-engineered multi-mutant; Marks Lab had sequences generated from a natural sequence model with zero assay-labeled data as well as sequences generated from the same model with conditioning on the assay-labeled data. The *in vitro* round experimentation was performed by our experimental partner International Flavors and Fragrances (Leiden, Netherlands).

### Individual Property Measurements

For each submitted enzyme variant, where feasible, two sequence-verified clones were assayed (see Methods), and their resulting measurements were averaged to evaluate expression levels, specific activity, and thermostability (Fig. 4). Comparative data between these replicates were used to assess assay noise and were found to have satisfactory reproducibility across all measured properties (Supp. Disc. Data Quality). For thermostability, the enzyme variants from teams SergiR1996 and MediumBio demonstrated superior performance relative to other groups. For specific activity, the primary property of interest, TUM emerged as the clear winner. Most teams showed comparable levels of expression; however, due to technical issues, the expression levels of the control protein were not directly comparable to those of the submitted variants. The control molecule’s expression was approximately estimated to be three times higher than actual, leading us to adjust the comparison to 30% of the control’s expression levels for fairness.

**Figure 4:**
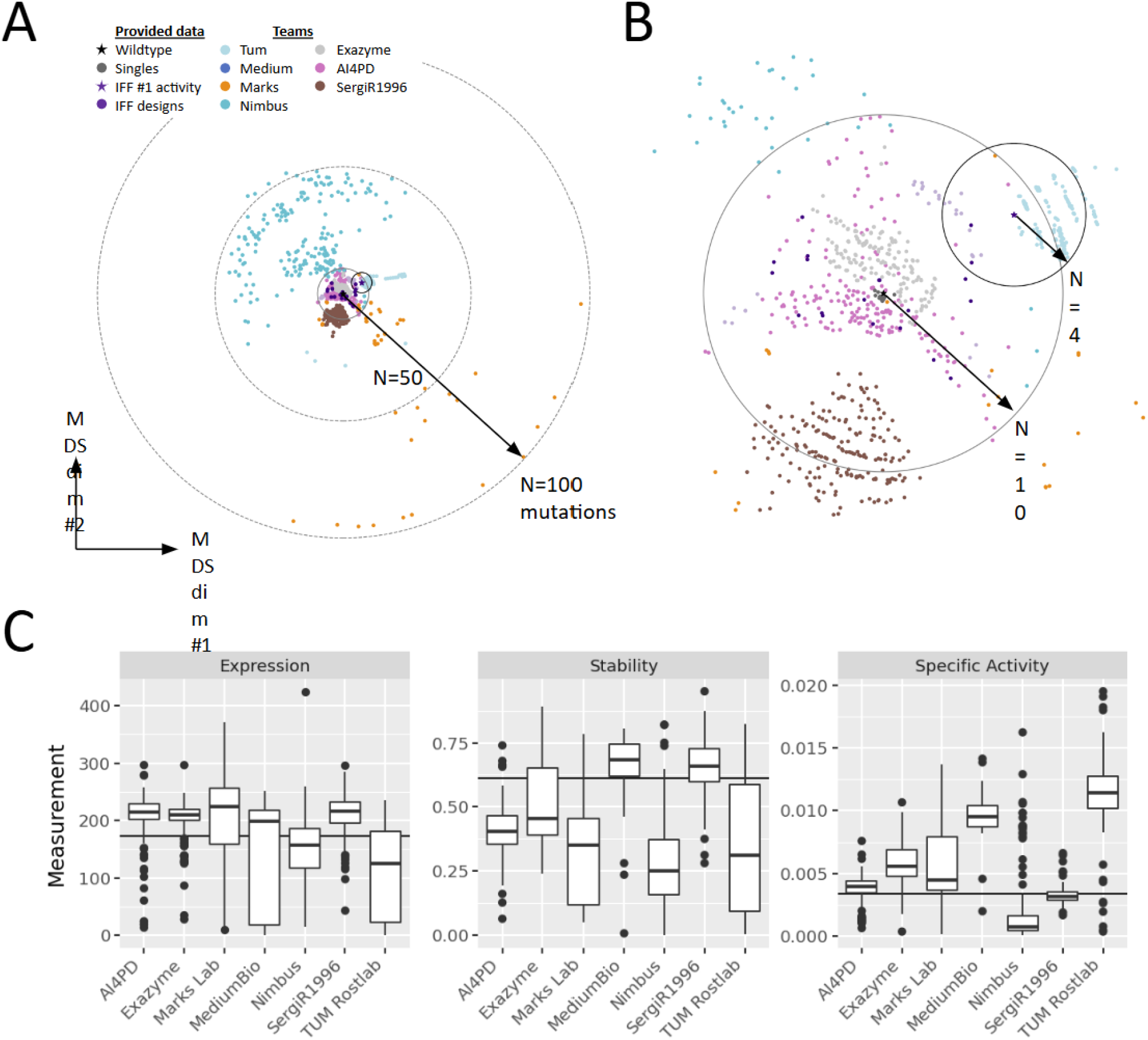
Individual Property Measurements. **(A)** Teams took diverse approaches, introducing as many as 100 mutations simultaneously, **(B)** and working from either the wildtype alpha-amylase (black star) or from the highest activity sequence among the provided combinatorial design data (purple star). **(C)** Variants for each team were measured for expression, stability, and specific activity, with the average value indicated. The level of the reference wild-type enzyme is indicated by the solid black line, except in the Expression plot where it indicates 30% of the wild-type enzyme. The units for Expression are in parts per million (ppm), Stability is unitless, and Specific Activity is in optical density per parts per million (OD / ppm).

### Final Evaluation of Designed Sequences

While all teams successfully engineered variants with improved activities, TUM Rostlab distinguished itself with both the top-scoring individual variant and the highest median performance among the qualifying variants that met the combined design criteria for expression and thermostability (Fig. 5). MediumBio and Marks Lab were close contenders, finishing second and third, respectively. Detailed numerical results for the variants that passed the design criteria are provided, including data on the best single variant activity and median activity across variants for each team (Supp. Table 4). Additionally, special recognition was given to teams for outstanding achievements in individual property improvements, combinations of these properties, and for the team with the greatest number of variants meeting the design criteria (Supp. Table 5).

**Figure 5:**
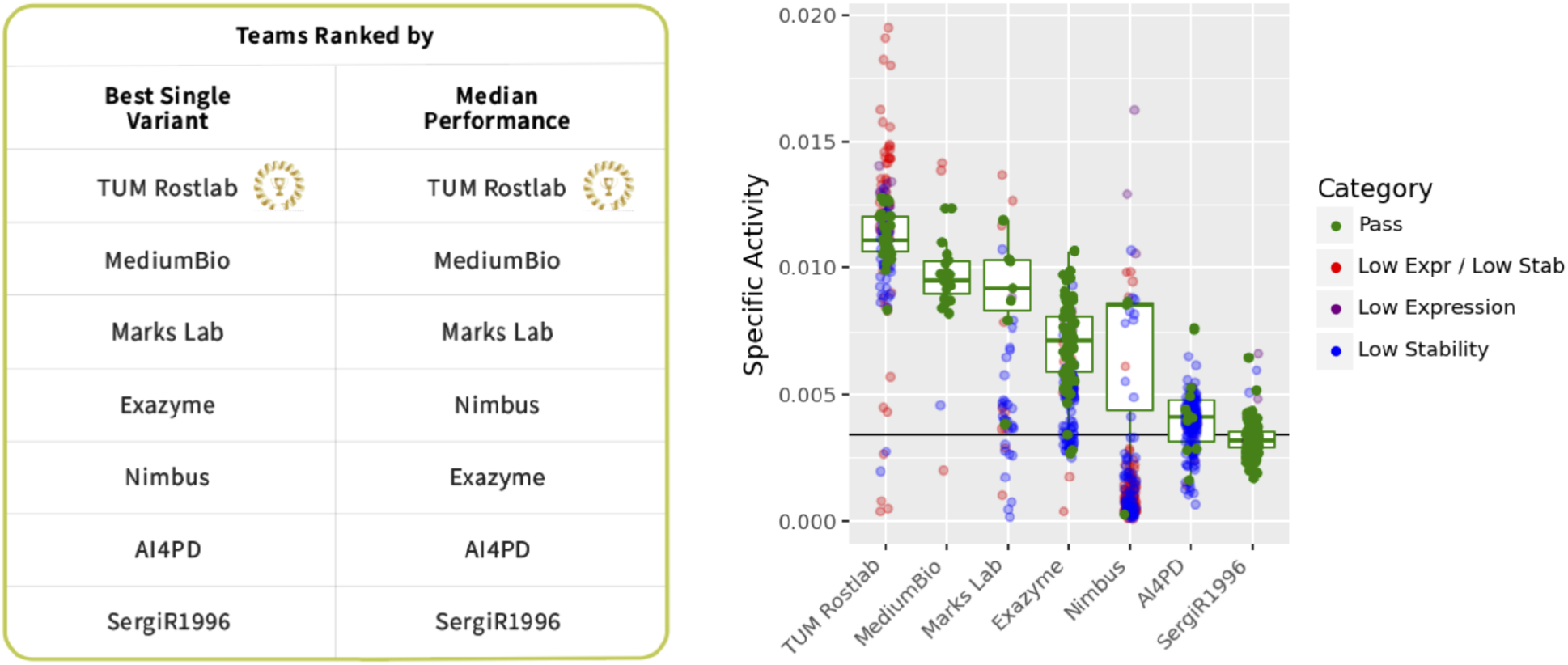
Final results for the Tournament. The level of the reference wild-type enzyme is indicated by the solid black line, except in the Expression plot where it indicates 30% of the wild-type enzyme. The units for specific activity are in optical density per parts per million (OD / ppm).

## Discussion

The Protein Engineering Tournament has introduced an innovative, community-driven platform to accelerate the advancement of computational protein engineering. This open science initiative combines *in silico* datasets for predictive modeling and in vitro experimentation for generative protein design. In creating a unified benchmark, the Tournament offers a communal yardstick for evaluating the performance of computational approaches in modeling and engineering protein function. The pilot tournament demonstrated these principles by offering never-before-seen datasets on protein function, novel mappings between computational approaches and experimental outcomes, and ample opportunities for community engagement and knowledge sharing.

The pilot Tournament’s *in vitro* round successfully produced a new **α**-amylase dataset including expression, thermostability, and specific activity measurements of ∼1,200 variants from the external partnership with International Flavors and Fragrances. In future iterations, the Tournament’s flexible structure allows for additional methods of generating in vitro data, including partnerships with: 1) external collaborators in academia or industry, 2) companies with Laboratory Data as a Service models, or 3) automation-enabled facilities known as cloud science laboratories. We expect our Tournaments to use a mixture of these methods to leverage their unique benefits and continually adapt to an ever-changing experimental ecosystem (Supp. Table 6).

The Tournament’s *in vitro* round also offers a unique setting for benchmarking across many types of computational models. The 2023 Protein Engineering Tournament showcased a broad spectrum of computational approaches, reflecting the diverse landscape of modern protein engineering.

Participants employed a range of techniques, from sequence- and structure-based models to protein large language models (LLMs) in order to perform this multi-objective optimization. TUM Rostlab and AI4PD utilized fine-tuning of protein language models, and TUM Rostlab further built upon these large-scale models using reinforcement training. MediumBio applied a combination of greedy recombination of beneficial mutations and model-guided protein sequence space sampling.

Alternatively, Nimbus employed structural modeling tools like PyRosetta^12^ and AlphaFold^13^. Marks Lab used natural sequence models to sample some sequences zero-shot and some sequences via assay-labeled conditional generation, and AI4PD also fine-tuned language models, SergiR1996 used a structure based and genetic algorithms to perform multi-objective optimization. Notably, the breadth of techniques implemented by top teams highlights the lack of consensus around the ‘best’ method for protein design. The choice to build on top of IFF’s prior engineering, and the number of mutations introduced (Figure 4A-B), strongly stratified team performance. For instance, the top activity designs were within 1-4 mutations of the top activity design discovered by IFF’s engineering efforts. By contrast, the average design distance was 15 mutations and the most distant examples exceeded 100 mutations. Though it is tempting to draw conclusions about the different strengths of the computational models each team invokes, significant high-level differences in approach may have made the largest impact on the design outcomes. Future Tournaments will enable teams to have multiple avenues to participate in the in vitro round, to maximize the diversity of computational approaches and number of teams that can participate (Supp. Disc. Participation).

In future tournaments the design challenges will also expand to encompass more domains of function (Supp. Fig. 3). The order in which we introduce new functions will be driven by practical application, technical feasibility, and amenability to high-throughput experimentation. As our computational methods improve, our challenges will expand into increasingly more difficult and complex domains, such that the frontier of scientific capabilities is always represented in the Tournament’s challenges. While the first pilot Tournament solely focused on enzyme engineering, we see enzyme engineering and protein binder design as strong initial candidates for the first official Tournament in 2025. As we look ahead to the first official Tournament, we anticipate that this initiative will contribute significantly to the evolving landscape of protein engineering and enable the scientific community to conquer the grand challenge of protein design.

## Methods

All data, metadata, team submissions, analysis scripts, figures, and team abstracts are available on our Github: https://github.com/the-protein-engineering-tournament/pet-pilot-2023. This resource offers an in-depth look at the diverse strategies employed during the tournament, serving as a valuable asset for researchers and enthusiasts in the field.

### *in silico* Analysis Metrics and Scoring

For all *in silico* events, excluding Xylanase, Spearman correlation was chosen as the evaluation metric to compare submitted predicted properties to their measured values. The choice of Spearman correlation was driven by several reasons. First, Spearman correlation is robust to non-linear relationships and does not assume linearity. As a rank-based metric, it is less sensitive to outliers compared to Pearson correlation, which is especially beneficial in the zero shot setting where the ranges of submission were arbitrary. Additionally, its ease of interpretation and widespread use in the protein engineering literature supported this choice.

The only event not analyzed by Spearman correlation was the Xylanase supervised learning event. The Xylanase event was analyzed using a weighted F1 score because it was a classification task (three options: No Expression, Low Expression, and Good Expression). In multi-modal events (i.e., events asking teams to predict more than one property) Spearman correlation across all properties in an event were averaged together, resulting in a final averaged score that allowed us to rank teams from highest to lowest.

Teams were awarded points based on their rank for each individual event using a reverse-rank reward system (e.g., teams coming in 1st, 2nd, and 3rd place were awarded 3, 2, and 1 points respectively). This system rewarded teams for doing well in events and for competing in multiple events, while normalizing to the number of competitors per event. In the event of a tie, both teams were awarded the same amount of points, sharing the higher rank.

### *in vitro* Experimental Methods

Participants’ sequences were cloned and sequence verified. When possible, two clones for each variant were characterized in duplicate for three properties: expression, thermostability, and specific activity. All experimentation was performed by our automation partners, International Flavors and Fragrances, following substantially the same methods that created the original **α**-amylase dataset provided to the teams.

### *in vitro* Expression Measurements

For the expression measurement, protein concentration in sample supernatants was determined using size exclusion chromatography (SEC). Samples were obtained by filtering broths from cultures grown in microtiter plates (MTPs) for 3 days at 37°C with shaking at 250 rpm and humidified aeration. Filtered samples were diluted and analyzed on HPLC (high pressure liquid chromatography) using an Acquity BEH300 200A SEC column (Waters Corp), with eluent containing 25 mM Acetate-buffer, pH 5.8 and 100 mM NaCl at a flow of 0.3 mL/min. Absorbance was measured at 220 nm and integrated peak area was determined. Sample concentrations were calculated by comparison to the purified **α**-amylase standard of known concentration used as a calibrator.

### *in vitro* Specific Activity and Thermostability Measurements

A starch hydrolysis assay was used to determine specific activity and thermostability of **α**-amylase variants. First, 1% (w/v) corn starch (Ingredion Inc.) was dissolved in dilution buffer (50 mM MOPS buffer pH 7.15 with 5 ppm (parts per million) Ca^2+^ and 20 ppm Na^+^ with 0.002% TWEEN-80®). Dissolved corn starch was boiled in a microwave, then cooled to room temperature overnight with gentle stirring. Enzyme aliquots (appr. 0.05 μg/L) were added to 1% corn starch (50 μL final volume) in a 384-well polypropylene v-bottom microtiter plate (Greiner 384w PP MTP, Catalog No. 781280) and mixed on a BioShake 3000 plate mixer (QInstruments) for 15 seconds at 2800 rpm. Reactions were sealed with a heat seal for 1.5 seconds at 180°C and incubated in an iEMS shaking incubator (Thermo Scientific) for 30 min at 60°C with 1150 rpm shaking. After incubation, the plates were cooled down at room temperature and spun down briefly to remove condensate on the seals. The bicinchoninic acid (BCA) assay used for detection of reducing ends was carried out using Pierce BCA Assay Kit (Catalog number 23225, ThermoFisher Scientific). After the starch hydrolysis reactions were cooled to room temperature, 5 μL of the reaction mixture were added to 30 μL BCA reagent solution in a 384-well PCR plate (Eppendorf twin.tec PCR MTP 384, Catalog No. 951020737). This mixture was heated to 95°C for 3 minutes, cooled to room temperature, and 25 μL of this BCA reaction were then transferred to a 384-well polystyrene flat-bottom read plate (Greiner 384-well PS MTP, Catalog No. 781101) and absorbance was measured at 562 nm.

A stress incubation was performed on the enzyme samples at 85C for 20 minutes and the starch hydrolysis assay was run as described above to determine activity after the stress incubation. Reducing ends were detected using BCA assay by measuring absorbance at 562 nm.

Activity was calculated from the number of reducing ends generated by enzymatic breakdown of corn starch, as quantified by the BCA assay. Specific activity was then calculated by dividing the activity (measured before the stress incubation) by the protein concentration value previously determined by the expression measurement. Stability was calculated as the ratio of activity after the stress incubation to before the stress incubation (i.e., residual activity).

### *in vitro* Sequence Design Analysis

To assess which variants met all predetermined design criteria, each variant was initially classified based on its expression and stability: categorized as having ‘too low expression’, ‘too low stability’, ‘both low stability and expression’, or ‘pass’ for those meeting all criteria (Supp. Table 3). The variants that passed were further analyzed to identify the team that developed the variant with the highest specific activity (Supp. Table 4).

## Acknowledgements

The Protein Engineering Tournament is operated by Align to Innovate, a non-profit organization dedicated to improving the reproducibility, scalability, and shareability of life science research through community-driven initiatives. Align to Innovate receives philanthropic funding in part from Griffin Catalyst. The Tournament is run by a planning committee composed of Align to Innovate employees and volunteers from the protein engineering community.

The Tournament Planning Committee thanks all dataset donors for helping to make the pilot Tournament possible. We thank the Polly Fordyce Group, Stanford University, and specifically Dr. Craig Markin and Prof. Polly Fordyce for contributing an Alkaline Phosphatase PafA dataset. We thank Prof. Dr. U.T. Bornscheuer and M.Sc. M. J. Menke from the University of Greifswald, Dept. of Biotechnology & Enzyme Catalysis for contributing a dataset on activity data of transaminase-catalyzed conversion of bulky substrates. We thank Codexis for contributing an Imine Reductase dataset. We thank International Flavors and Fragrances (IFF), the Sarel Fleishman Lab from the Weizmann Institute of Science for contributing the Xylanase dataset. We thank the students and faculty members of the Design to Data (D2D) undergraduate research program, led by Ashley Vater and Professor Justin B. Siegel at University of California, Davis.

We would also like to thank our *in vitro* round experimental partners at IFF’s R&D facilities in Leiden, Netherlands. The in vitro libraries were cloned by Harm Mulder and Rei Otsuka. HPLC based concentration determination was performed by Laurens Lammerts. Biochemical activity and stability characterization was performed by Sina Pricelius, Lydia Dankmeyer, Veli Alkan, Viktor Alekseyev, and Frits Goedegebuur.

