## Supplemental Materials for "Results of the Protein Engineering Tournament: An Open Science Benchmark for Protein Modeling and Design"

### Supplemental Figures and Tables

Supplemental Table 1: **Participating teams from each round of the Tournament.**

| ***in silico* teams** | ***in vitro* teams** |
|---|---|
| Arnold Lab | AI4PD |
| Exazyme | Exazyme |
| Marks Lab | Marks Lab |
| Nimbus | Medium Bio |
| ProtEng4All | Nimbus |
| SergiR1996 | SergiR1996 |
| TUM Rostlab | TUM Rostlab |

Supplemental Table 2: **Suggested citations, links to GitHub, and additional references material for each dataset.** Preferred citations and optional additional references are shown in their original form from Dataset donors. The GitHub entries link directly to the relevant folders in the Protein Engineering Tournament's 2023 pilot repository.

| **Enzyme Dataset** | **Preferred citation** | **GitHub Link** | **Additional documentation and resources** |
|---|---|---|---|
| Aminotransferase | This dataset is not published (yet) in any scientific journal. Please use the citation: "Menke, M. J.; Bornscheuer, U. T. Activity data of quadruple mutants of 3FCR_4M transaminase for bulky substrates. 2023" | https://github.com/the-protein-engineering-tournament/pet-pilot-2023/tree/main/in_silico_zero-shot/input/Aminotransferase%20(in%20silico_%20Zero-Shot) | • Transaminase engineering based on related data: „Ao, Y.-F.; Pei, S.; Xiang, C.; Menke, M. J.; Shen, L.; Sun, C.; Dörr, M.; Born, S.; Höhne, M.; Bornscheuer, U. T. Structure- and data-driven protein engineering of transaminases for improving activity and stereoselectivity. Angew. Chem. Int. Ed. 2023, 62, e20230166“.<br>• The 3DM tool to plan the mutations: „Kuipers, R.K.P., Joosten, H.-J., Verwiel, E., Paans, S., Akerboom, J., van der Oost, J., Leferink, N.G.H., van Berkel, W.J.H., Vriend, G. and Schaap, P.J. (2009), Correlated mutation analyses on super-family alignments reveal functionally |

| | | | |
|---|---|---|---|
| | | | important residues. Proteins, 76: 608-616. https://doi.org/10.1002/prot.22374 “ |
| α-Amylase | F. van der Flier, D. Estell, S. Pricelius, L. Dankmeyer, S. van Stigt Thans, H. Mulder, R. Otsuka, F. Goedegebuur, L. Lammerts, D. Staphorst, A.D.J. van Dijk, D. de Ridder and H. Redestig. Preprint bioRxiv 2023. https://www.biorxiv.org/content/10.1101/2023.09.25.559319v1 | *in silico* supervised track: https://github.com/the-protein-engineering-tournament/pet-pilot-2023/tree/main/in_silico_supervised/input/Alpha-Amylase%20(In%20Silico_%20Supervised)<br><br>*in silico* zero-shot track: https://github.com/the-protein-engineering-tournament/pet-pilot-2023/tree/main/in_silico_zero-shot/input/Alpha-Amylase%20(in%20silico_%20Zero-Shot)<br><br>*In vitro* round: https://github.com/the-protein-engineering-tournament/pet-pilot-2023/tree/main/in_vitro/input | The AmyE training data for the in silico challenge was generated as part of published patents 1-4 and further disseminated in 5.<br><br>1. L. G. Cascao-Pereira, R. Chin, W. A. Cuevas, D. A. Estell, S.-K. Lee, M. J. Pepsin, S. D. Power, S. W. Ramer, C. A. Requadt, A. Shaw, A. R. Toppozada, and L. Wallace, “Uses of an α-amylase from bacillus subtilis.” European Patent Application, Ep 2.698,434 (A1), Aug 2014.<br>2. L. G. Cascao-Pereira, W. A. Cuevas, D. A. Estell, S.-K. Lee, S. D. Power, S. W. Ramer, A. Toppozada, and L. Wallace, “Variant α-amylases from bacillus subtilis and methods of uses thereof.” US Patent, US 8,323,945 (B2), Dec 2012.<br>3. L. G. Cascao-Pereira, W. A. Cuevas, D. A. Estell, S.-K. Lee, S. D. Power, S. W. Ramer, A. Toppozada, and L. Wallace, “Variant α-amylases from bacillus subtilis and methods of uses thereof.” US Patent, US 8,975,056 (B2), Mar 2015.<br>4. W. A. Cuevas, S.-K. Lee, S. W. Ramer, A. Shaw, A. R. Toppozada, D. E. Estell, L. Wallace, R. Chin, C. A. Requadt, S. D. Power, and M. J. Pepsin, “Variant α-amylases from bacillus subtilis and methods of use thereof.” US Patent, US 9,090,887 (B2), Jul 2015.<br>5. F. van der Flier, D. Estell, S. Pricelius, L. Dankmeyer, S. van Stigt |

| | | | |
|---|---|---|---|
| | | | Thans, H. Mulder, R. Otsuka, F. Goedegebuur, L. Lammerts, D. Staphorst, A.D.J. van Dijk, D. de Ridder and H. Redestig. Preprint bioRxiv 2023. https://www.biorxiv.org/content/10.1101/2023.09.25.559319v2 |
| Alkaline phosphatase PafA | Markin, C. J., Mokhtari, D. A., Sunden, F., Appel, M. J., Akiva, E., Longwell, S. A., Sabatti, C., Herschlag, D. & Fordyce, P. M. Revealing enzyme functional architecture via high-throughput microfluidic enzyme kinetics. Science 373, eabf8761 (2021) | https://github.com/the-protein-engineering-tournament/pet-pilot-2023/tree/main/in_silico_supervised/input/Alkaline%20phosphatase%20PafA%20%20(In%20Silico_%20Supervised) | All data are available in a registered Open Science Foundation Repository (DOI: 10.17605/OSF.IO/QRN3C) |
| $\beta$- glucosidase B | D2D Database. 2018-. University of California, Davis: Design to Data. [updated 2024 Feb 15; accessed Apr 2023]. https://d2dcure.com/data/?protein=BglB | https://github.com/the-protein-engineering-tournament/pet-pilot-2023/tree/main/in_silico_supervised/input/beta-glucosidase%20B%20%20(In%20Silico_%20Supervised) | • Vater, A., Mayoral, J., Nunez-Castilla, J., Labonte, J. W., Briggs, L. A., Gray, J. J., ... & Siegel, J. B. (2020). Development of a broadly accessible, computationally guided biochemistry course-based undergraduate research experience. Journal of Chemical Education, 98(2), 400-409<br>• Huang, P., Chu, S. K., Frizzo, H. N., Connolly, M. P., Caster, R. W., & Siegel, J. B. (2020). Evaluating protein engineering thermostability prediction tools using an independently generated dataset. ACS omega, 5(12), 6487-6493.<br>• Carlin, D. A., Hapig-Ward, S., Chan, B. W., Damrau, N., Riley, M., Caster, R. W., ... & Siegel, J. B. (2017). |

| | | | |
|---|---|---|---|
| | | | Thermostability and kinetic constants for 129 variants of a family 1 glycoside hydrolase reveal that enzyme activity and stability can be separately designed. PLoS One, 12(5), e0176255. |
| Imine reductase | This dataset falls within the claims in the following patent application: Yi, Xiang, Oscar Alvizo, Ravi David Garcia, David Entwistle, Charlene Ching, Nandhitha Subramanian, and James Nicholas Riggins. 2020. ENGINEERED GLUCOSE DEHYDROGENASES AND METHODS FOR THE REDUCTIVE AMINATION OF KETONE AND AMINE COMPOUNDS. PCT/US2020/029517. | https://github.com/the-protein-engineering-tournament/pet-pilot-2023/tree/main/in_silico_supervised/input/Imine%20reductase%20(In%20Silico_%20Supervised) | |
| Xylanase | Listov, D.; Lipsh-Sokolik, R.; Rosset, S.; Yang, C.; Correia, B. E.; Fleishman, S. J. Assessing and Enhancing Foldability in Designed Proteins. *Protein Sci.* **2022**, *31* (9), e4400. | https://github.com/the-protein-engineering-tournament/pet-pilot-2023/tree/main/in_silico_zero-shot/input/Xylanase%20(in%20silico_%20Zero-Shot) | |

Supplemental Table 3: **Variant Sequence Statistics.** Number of variant sequences at each stage that were characterized for each team from submission to screening, further detailed by the number of sequences that passed all design criteria, had low expression, had low stability, or had both low expression and stability.

| | Number of sequences: | | | | | |
|---|---|---|---|---|---|---|
| Team | Submitted | Cloned & Screened | Passed Stability & Expression Criteria | Had Low Expression | Had Low Stability | Had Low Expression & Stability |
| TUM Rostlab | 200 | 199 | 30 | 18 | 38 | 54 |
| Marks Lab | 157 | 54 | 7 | 1 | 29 | 10 |
| MediumBio | 30 | 30 | 19 | 0 | 1 | 3 |
| Exazyme | 211 | 173 | 63 | 7 | 92 | 10 |
| Nimbus | 200 | 198 | 3 | 4 | 71 | 116 |
| SergiR1996 | 200 | 190 | 154 | 18 | 15 | 3 |
| AI4PD | 200 | 189 | 10 | 1 | 161 | 13 |

Supplemental Table 4: **Numerical values of specific activity for variants that passed the design criteria.** We show the numerical values for both the best single variant's specific activity and median specific activity across each team's variants.

| **Team** | **Best Single Variant Specific Activity (OD / ppm)** | **Median Specific Activity (OD / ppm)** |
|---|---|---|
| TUM Rostlab | 0.0128 | 0.0111 |
| MediumBio | 0.0123 | 0.0095 |
| Marks Lab | 0.0119 | 0.0092 |
| Exazyme | 0.0107 | 0.0071 |
| Nimbus | 0.0086 | 0.0085 |
| AI4PD | 0.0076 | 0.0041 |
| SergiR1996 | 0.0065 | 0.0032 |

Supplemental Table 5. **Honorable mentions.** Teams that had outstanding variants for each individual property measurements, as well as combinations of measurements were highlighted. This included the team with the highest number of variants passing all design criteria.

| Award | Team |
|---|---|
| Highest Specific Activity | TUM Rostlab |
| Highest Stability | SergiR1996 |
| Highest Expression | Nimbus |
| Highest Specific Activity & Stability | TUM Rostlab |
| Highest Expression & Stability | SergiR1996 |
| Highest Expression & Specific Activity | TUM Rostlab |
| Highest Specific Activity & Expression & Stability | TUM Rostlab |
| Most Variants Passing Design Criteria | SergiR1996 |

Supplemental Table 6: **Avenues for *in vitro* round data generation.** The Tournament allows for flexible selection of how *in vitro* round data is generated.

| Approach | Description | Benefits | Limitations |
|---|---|---|---|
| Experimental Partnerships | Form collaborations with experimental biology laboratories in academia or industry to conduct the experiments | • Can publish detailed descriptions of the assay protocols to facilitate experimental reproduction and ongoing benchmarking. | Participants without experimental capabilities may lack a clear path to continue benchmarking new computational methods after the |

| | | | |
|---|---|---|---|
| | | <ul><li>Leverages the expertise of experienced biologists to lead assay development and execution, improving the likelihood of high-quality experimental outcomes.</li></ul> | Tournament has concluded. |
| Laboratory Data as a Service | Utilizes an emerging business model known as , which enables researchers to order protein characterization assays directly online. Researchers select the desired assay, provide a list of protein sequences, and place their order. Upon completion of the experiments, the resultant data is uploaded for analysis. | <ul><li>Offers high feasibility for the Tournament and allows researchers without experimental capabilities to continue benchmarking beyond the Tournament.</li></ul> | Experimental details of these assays are often not made publicly available. This poses challenges for the experimental reproduction of the Tournament's results, as research groups may lack the necessary information and instrumentation necessary to do so. |
| Cloud Laboratories | Leverages automation-enabled facilities known as cloud science laboratories. Researchers utilize a symbolic laboratory programming language that specifies the instructions for each step of the assay protocol. Scientists submit their | <ul><li>Holds significant potential to improve the accessibility and reproducibility of life science research, enabling scientists to share and reproduce experimental protocols with the same ease as software today.</li><li>At the conclusion of the Tournament, the cloud laboratory protocols we developed for our assays</li></ul> | Tournament organizers would need to develop and validate these cloud-based assays, so this approach requires the most significant investment of time and effort. |

| | | |
|---|---|---|
| | protocols through a web portal; automated wet-lab robots then perform each experimental step and the resulting data is uploaded for analysis. | would be made openly available to the scientific community. This enables both computational and experimental researchers to access our assays, reproduce our results, and benchmark new computational methods.<br>● As each Tournament introduces new protein engineering challenges, this approach will lead to an ever-expanding corpus of open-source protein characterization assays to benchmark computational methods for years to come. |

### Supplementary Discussion: Data Quality

To account for experimental variation, selected clones for each variant were randomly designated as Replicate A and Replicate B in each property measurement. Comparative data between these replicates were used to assess assay noise (Supp. Fig. 1). The correlation between replicates was quantitatively assessed using Spearman's Rho, yielding coefficients of 0.79 for expression, 0.88 for thermostability, and 0.92 for specific activity. These values were found to be sufficient for the property measurements.

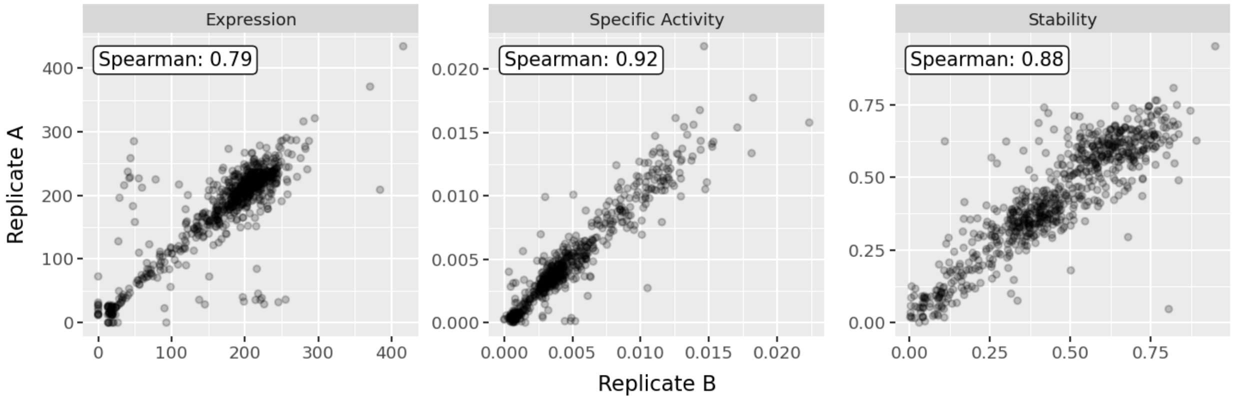

Supplemental Figure 1: **Data Quality.** When possible, variants were tested in replicate for each of the property measurements and compared against each other using Spearman's Rho to assess data quality.

#### Supplementary Discussion: Participation

The first round of the Protein Engineering Tournament, the *in silico* round, was open to any and all researchers interested in participating. Interested teams composed of one or more individuals were able to download challenge data and upload final predictions.

To allow the greatest number of teams to participate in the *in vitro* round, while being conscious of the costs associated with DNA synthesis and experimental characterization, we proposed two avenues for admission in the pilot Tournament: 1) high performance in the *in silico* round, 2) special invitation for generative teams. In future iterations, we propose an additional avenue: paid entry (Supp. Fig. 2). The first path focuses on rewarding innovative research teams who have demonstrated promising computational approaches in the *in silico* round. The second path is designed for groups with expertise in generative protein design but less experience in the *in silico* round's property prediction tasks. For the final path, a paid entry route will be available in future Tournaments, primarily aimed at well-funded corporate labs, to broaden participation and allow a wider range of methods to be evaluated in the tournament.

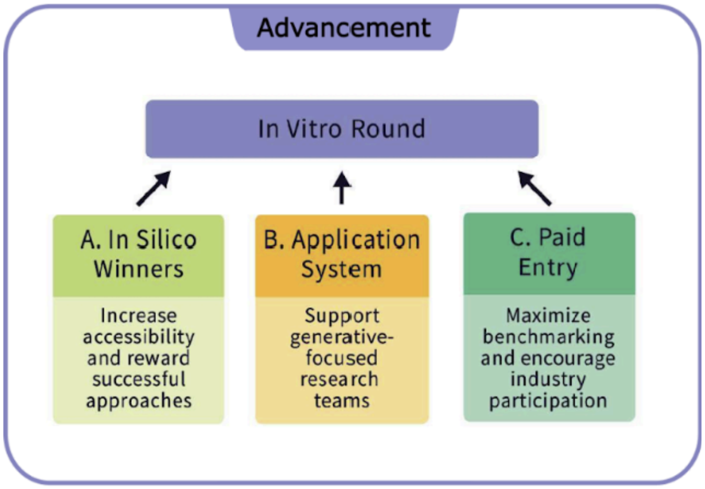

Supplementary Figure 2. **Avenues for in vitro participation.** There will be three avenues for participants to enter into the *in vitro* round, with each avenue catering to a unique audience.

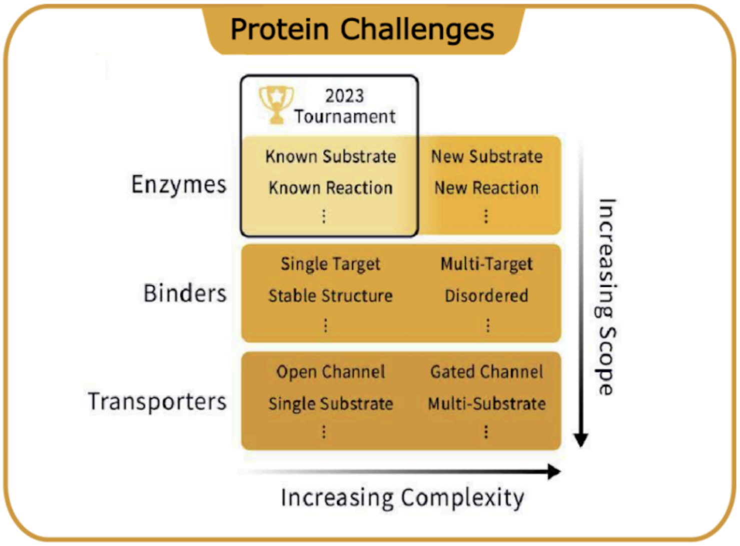

Supplemental Figure 3: **Selecting Protein Design Challenges.** The Tournament will continually expand to new domains of protein design; in each domain, we will continually select challenges that push the limits of current techniques. Our 2023 Pilot Tournament focused on Enzyme Design.
